# Vitamin C and Transferrin Reduce RNA Methylation in Mouse Embryonic Stem Cells

**DOI:** 10.1101/2023.02.23.529811

**Authors:** Gabrielle Brandt, Laura J. Sedivy, Matthew Mitchell, Christopher J. Phiel

## Abstract

Methylation of mRNA on adenosine bases (referred to as m^6^A) is the most common internal modification of mRNA in eukaryotic cells. Recent work has revealed a detailed view of the biological significance of m^6^A-modified mRNA, with a role in mRNA splicing, control of mRNA stability, and mRNA translation efficiency. Importantly, m^6^A is a reversible modification, and the primary enzymes responsible for methylating (Mettl3/Mettl14) and demethylating RNA (FTO/Alkbh5) have been identified. Given this reversibility, we are interested in understanding how m^6^A addition/removal is regulated. Recently, we identified glycogen synthase kinase-3 (Gsk-3) activity as a mediator of m^6^A regulation via controlling the levels of the FTO demethylase in mouse embryonic stem cells (ESCs), with Gsk-3 inhibitors and Gsk-3 knockout both leading to increased FTO protein and decreased m^6^A mRNA levels. To our knowledge, this remains one of the only mechanisms identified for the regulation of m^6^A modifications in ESCs. Several small molecules that have been shown to promote the retention of pluripotency of ESCs, and interestingly, many have connections to the regulation of FTO and m^6^A. Here we show that the combination of Vitamin C and transferrin potently reduces levels of m^6^A and promotes retention of pluripotency in mouse ESCs. Combining Vitamin C and transferrin should prove to be valuable in growing and maintaining pluripotent mouse ESCs.

## Introduction

Mouse embryonic stem cells (ESCs) are derived from blastocyst stage embryos and have the capacity to give rise to all three embryonic germ layers (ectoderm, mesoderm, and endoderm) and the cells derived from these germ layers [1]. This cellular plasticity, known as pluripotency, is one of the cardinal features, along with self-renewal, that makes ESCs unique from differentiated cell types [2]. Work from numerous labs has demonstrated that ESCs must be actively retained in a pluripotent state or they will spontaneously differentiate in culture [1, 3]. This requires the addition of media supplements such as leukemia inhibitory factor (LIF) to maintain pluripotency [4]. Other proteins and small molecules have also been shown to assist in promoting this pluripotent state, including small molecule inhibitors of glycogen synthase kinase-3 (Gsk-3) [5] and mitogen-activated kinase kinase (MEK) [6], ascorbic acid [7], and α-ketoglutarate [8], to name a few. There are two main pluripotent states for mouse ESCs in culture, referred to as naïve and primed [9]. Naïve ESCs most closely resemble *in vivo* ESCs found in early blastocyst embryos (E4.5). Primed ESCs represent ESCs that have begun the progression toward a more differentiated cell, more specifically resembling ESCs derived from slightly older, post-implantation embryos *in vivo* (E6.5), and are also referred to as EpiSCs (epiblast stem cells) [10]. Mouse ESCs grown in the presence of LIF are essentially primed ESCs, while the combination of Gsk-3 and MEK inhibitors have been shown to keep ESCs in a naïve state [6]. Therefore, rather than existing in a single state, ESCs are on a continuum, with naïve ESCs representing completely undifferentiated cells, while primed ESCs are in a state between naïve and differentiated cells. Differences in ESC pluripotency can be identified by measuring the mRNA abundance of marker genes [11]. For example, *Nanog* is highly expressed in all ESCs [12, 13], while *Fgf5* is a specific marker of primed ESCs [14]. Another means of assessing pluripotency is by quantifying the relative amount of RNA methylation [15, 16]. Finally, another reliable way to determine pluripotency is by cellular phenotype visualized by microscopy. Primed ESCs tend to grow in flat clusters as a monolayer, while naïve ESCs grow in distinct rounded spheres composed of tightly packed ESCs [9].

There are over 100 different chemical modifications that are found on RNA molecules, and the biological role of many of these modifications is unknown. One modification, methylation of adenosine bases at the C6 position (referred to as m^6^A), was discovered almost 50 years ago, and is the most common internal (non-cap) mRNA modification [17–19]. Data indicating the functional importance of m^6^A bases in mRNA was difficult to obtain at that time, and only recently have those functions begun to emerge [20–22]. Enrichment of m^6^A followed by next-generation sequencing (m^6^A-seq) has revealed a great deal of information about the precise mRNAs that contain m^6^A, as well as their abundance and distribution along mRNAs [23, 24]. Importantly, m^6^A is a reversible modification, and enzymes have been identified that facilitate the addition and removal of the m^6^A modifications (Mettl3, and FTO and Alkbh5, respectively) [25–27]. In addition, reader proteins have been discovered whose function is to recognize and bind to mRNA modified with m^6^A (YTHDF, IGF2BP) [28, 29]. The discovery of proteins that control m^6^A mRNA levels has allowed for studies to determine the biological consequences of enhancing or depleting m^6^A tags on mRNA. It has recently been reported that YTHDF proteins and IGF2BP proteins have essentially opposite functions with respect to the effects on mRNA modified by m^6^A; YTHDF2 binding to m^6^A promotes mRNA decay [28], while IGF2BP binding stabilizes m^6^A modified mRNAs [30]. In mouse ESCs, FTO is the main RNA demethylase since Alkbh5 is expressed almost exclusively in testes.

We have previously identified a novel role for Gsk-3 in promoting ESC pluripotency by regulating m^6^A RNA abundance via the RNA demethylase FTO [31]. Under normal conditions, Gsk-3 phosphorylates FTO, tagging FTO for ubiquitination and degradation by the proteasome. Thus, FTO levels are kept low and levels of m^6^A are relatively higher. When Gsk-3 is inhibited by small molecules or genetically deleted in ESCs, FTO protein levels accumulate, leading to a relative reduction in m^6^A. Since Gsk-3 inhibitors are used to promote ESC pluripotency, and Gsk-3 DKO ESCs are unable to differentiate, the direct connection between Gsk-3, FTO, and m^6^A levels is intriguing and likely hints at important mechanisms with respect to the regulation of the pluripotent state in ESCs. Several small molecules have been implicated in promoting the retention of mouse ESCs in culture. Interestingly, several of the same molecules have also been shown to regulate FTO function in various settings. In this study, we have examined these molecules not just for their effects on pluripotency, but also on mRNA methylation.

## Results and Discussion

### Wild-type ESCs

We treated WT ESCs with α-ketoglutarate, NADPH, Vitamin C and transferrin, and in some cases combined these treatments. Each treatment period was for 48 hours, after which time cells were collected and RNA was extracted and quantified. α-ketoglutarate has been shown to enhance the retention of ESC pluripotency. Interestingly, FTO is an α-ketoglutarate-dependent enzyme, suggesting that the effect of α-ketoglutarate may be via enhancing the function of FTO. Similarly, the cellular metabolite NADPH was recently shown to be a potent co-factor for FTO. Vitamin C, a known promoter of ESC pluripotency, has also been implicated as a co-factor for FTO. And since FTO also requires iron (specifically Fe^+2^), we investigated a role for transferrin, the receptor that transports iron into cells. Other conditions, such as ESCs grown in the absence of LIF, were included as controls. We then subjected the RNA we isolated to RTqPCR for to measure the abundance of *Nanog* and *Fgf5*, as well as m^6^A ELISA to quantify levels of mRNA methylation.

Individually, some treatments resulted in reduced m^6^A levels. For example, as seen in Table 1, NADPH resulted in a modest 5.8% reduction in m^6^A, while transferrin and Vitamin C each reduced m^6^A levels more dramatically (19.5% and 25.2%, respectively). However, the combined treatment of Vitamin C and transferrin resulted in a 49% reduction in m^6^A levels. The changes in the Nanog:Fgf5 ratio trended in the same direction, with increased *Nanog:Fgf5* in ESCs with reduced m^6^A, although the correlation was not direct. For example, while Vitamin C treatment alone had a *Nanog:Fgf5* ratio of 19.2, the Vitamin C + transferrin treatment that had the lowest m^6^A values only had a *Nanog:Fgf5* ratio of 5.9. However, growing ESCs in the absence of LIF showed both the highest levels of m^6^A and lowest *Nanog:Fgf5* ratio, which was predicted. It was also unexpected to find that α-ketoglutarate had essentially no pluripotency-promoting effects. The levels of m^6^A were unchanged from untreated ESCs, while the *Nanog:Fgf5* ratio was 0.5. This shows us that gene expression levels of markers for different pluripotent states does not always correlate with m^6^A levels, despite a trend in the same direction.

**Table 1.**
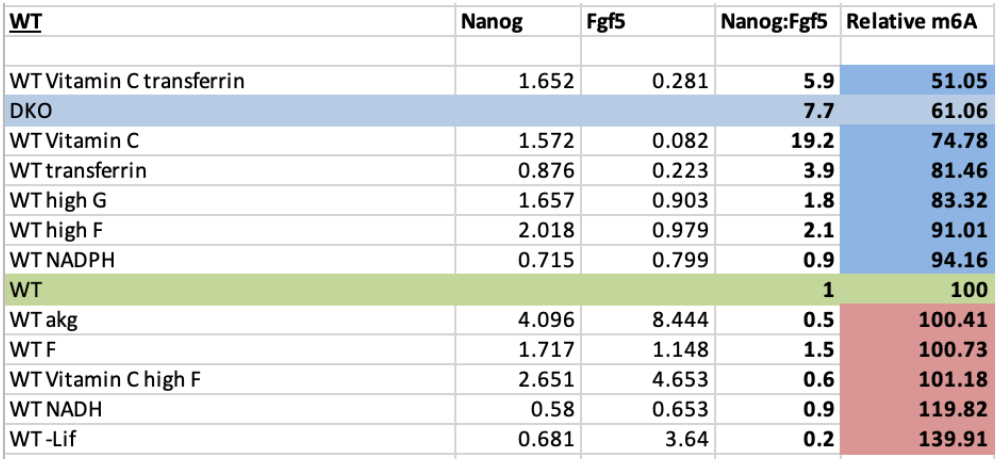

### Wild-type ESCs overexpressing FTO

Next, we assessed the effects of various small molecules in the presence of overexpressed FTO to see if the potency of the small molecules was enhanced. This was also done to confirm the effects were through FTO RNA demethylase activity. First, the overexpression of FTO alone dramatically reduced levels of m^6^A (53.6% reduction), while also showing an modest *Nanog:Fgf5* ratio of 2.8 (Table 2). There were some notable similarities and differences to the effects seen in WT ESCs that did not overexpress FTO. For example, Vitamin C treatment alone, despite have a high *Nanog:Fgf5* ratio of 14.0, actually displayed an increase in m^6^A levels. On the other hand, transferrin treatment resulted in both increased *Nanog:Fgf5* ratio (9.0) and led to a 23.5% reduction in m^6^A, which was even greater than in WT ESCs. However, the greatest effect on m^6^A levels was once again the combined treatment of Vitamin C and transferrin, with a remarkable 79.2% reduction in m^6^A abundance, as well as a *Nanog:Fgf5* ratio of 5.5. Taken together, these data strongly suggest that the combination of increased FTO expression plus Vitamin C and transferrin treatment is a potent modulator of m^6^A in mouse ESCs, and based on marker gene expression, promotes the retention of ESCs in a more naïve state.

**Table 2.**

| **WT w/FTO overexpressed** | **Nanog** | **Fgf5** | **Nanog:Fgf5** | **Relative m6A** |
| --- | --- | --- | --- | --- |
| WT FTO Vitamin C transferrin | 2.043 | 0.371 | **5.5** | **20.81** |
| WT FTO NADH | 2.083 | 0.375 | **5.6** | **40.96** |
| **WT FTO** | **3.012** | **1.074** | **2.8** | **46.39** |
| WT FTO Vitamin C akg | 1.566 | 0.224 | **7.0** | **47.76** |
| WT FTO akg NADH | 2.325 | 0.54 | **4.3** | **51.78** |
| WT FTO Vitamin C high F | 23.584 | 14.531 | **1.6** | **52.67** |
| WT FTO akg Gsk3i | 5.075 | 1.971 | **2.6** | **53.35** |
| WT FTO NADPH | 2.35 | 0.37 | **6.4** | **54.61** |
| **DKO** |  |  | **7.7** | **60.25** |
| WT FTO akg meki Gsk3i | 5.693 | 0.298 | **19.1** | **61.59** |
| WT FTO Vitamin C akg NADPH | 2.973 | 0.228 | **13.0** | **67.82** |
| WT FTO transferrin | 3.466 | 0.0384 | **9.0** | **76.47** |
| WT FTO high F | 3.223 | 10.746 | **0.3** | **77.71** |
| WT FTO Gsk3i | 6.001 | 14.701 | **0.4** | **78.93** |
| WT FTO Vitamin C transferrin NADPH |  |  |  | **81.55** |
| WT FTO Vitamin C akg NADH | 3.263 | 0.314 | **10.4** | **85.15** |
| WT FTO akg meki | 6.837 | 1.217 | **5.6** | **86.53** |
| WT FTO Vitamin C NADH | 2.659 | 0.272 | **9.8** | **90.4** |
| **WT** |  |  | **1.0** | **100.00** |
| WT FTO akg NADPH | 3.899 | 0.563 | **6.9** | **100.11** |
| WT FTO Vitamin C NADPH | 2.73 | 0.247 | **11.1** | **110.84** |
| WT FTO Vitamin C | 12.5 | 0.892 | **14.0** | **118.02** |
| WT FTO Vitamin C transferrin NADH |  |  |  | **144.66** |
| WT FTO akg | 7.44 | 21.565 | **0.3** | **146.34** |

### *Gsk-3* DKO ESCs

Previous work from our lab has shown that Gsk-3 activity is a key regulator of FTO protein stability and subsequent levels of m^6^A. Therefore, we treated *Gsk-3* DKO ESCs with the same small molecules to see if the effects seen in *Gsk-3* DKO ESCs could be enhanced further. Since FTO levels are higher in *Gsk-3* DKO ESCs, we were curious to test whether predicted co-factors for FTO would further reduce m^6^A levels and perhaps enhance retention of pluripotency. As expected based on the results in WT ESCs, treatment with Vitamin C and transferrin individually led to a decrease in m^6^A levels (23.1% and 27.4%, respectively), while also elevating the *Nanog:Fgf5* ratio (Table 3). Furthermore, the combination of Vitamin C and transferrin potently reduced m^6^A levels by 55.5%, while increasing the relative *Nanog:Fgf5* ratio by 9.0 compared to untreated *Gsk-3* DKO ESCs, which was the greatest effect on the pluripotency markers of any treatment. Therefore, in *Gsk-3* DKO ESCs, the combination of Vitamin C and transferrin was the most potent effect on both m^6^A reduction and *Nanog:Fgf5* ratio.

**Table 3.**

| **<u>DKO</u>** | **Nanog** | **Fgf5** | **Nanog:Fgf5** | **Relative m6A** |
| --- | --- | --- | --- | --- |
| DKO Vitamin C transferrin | 2.818 | 0.314 | **9.0** | **44.45** |
| DKO Vitamin C transferrin entacapone | 1.643 | 1.832 | **0.9** | **55.85** |
| DKO transferrin | 1.031 | 0.78 | **1.3** | **72.61** |
| DKO high F | 1 | 0.54 | **1.9** | **83.18** |
| DKO Vitamin C high F | 0.71 | 0.11 | **6.5** | **92.08** |
| DKO Vitamin C_New | 0.828 | 1.145 | **0.7** | **94.88** |
| **DKO** |  |  | **1** | **100.00** |
| DKO F | 1.06 | 1.03 | **1.0** | **115.17** |
| DKO No G | 1.24 | 59.92 | **0.0** | **117.86** |
| DKO akg meki | 1.57 | 0.23 | **6.8** | **127.28** |
| DKO -Lif | 0.66 | 1.1 | **0.6** | **137.82** |
| DKO akg | 1.08 | 1.17 | **0.9** | **161.06** |
| DKO high G | 2.88 | 2.76 | **1.0** | **254.78** |

### *Gsk-3* DKO ESCs overexpressing FTO

Finally, we wanted to see if overexpression of FTO in Gsk-3 DKO ESCs had an enhanced effect on mRNA methylation reduction without FTO overexpression. While overexpression of FTO did result in a reduction in m^6^A levels, the effect was modest (19.2%) and was accompanied by a similarly modest increase in *Nanog:Fgf5* ratio (2.0) (Table 4). In the scenario, similar to overexpression of FTO in WT ESCs, Vitamin C treatment had no effect on m^6^A levels despite showing an increase in *Nanog:Fgf5* ratio (4.1 compared to untransfected *Gsk-3* DKO ESCs. Transferrin treatment also had essentially no effect on m^6^A levels. However, the combined treatment of Vitamin C and transferrin once again was a potent effector of m^6^A levels, reducing m^6^A by 41.6%.

**Table 4.**
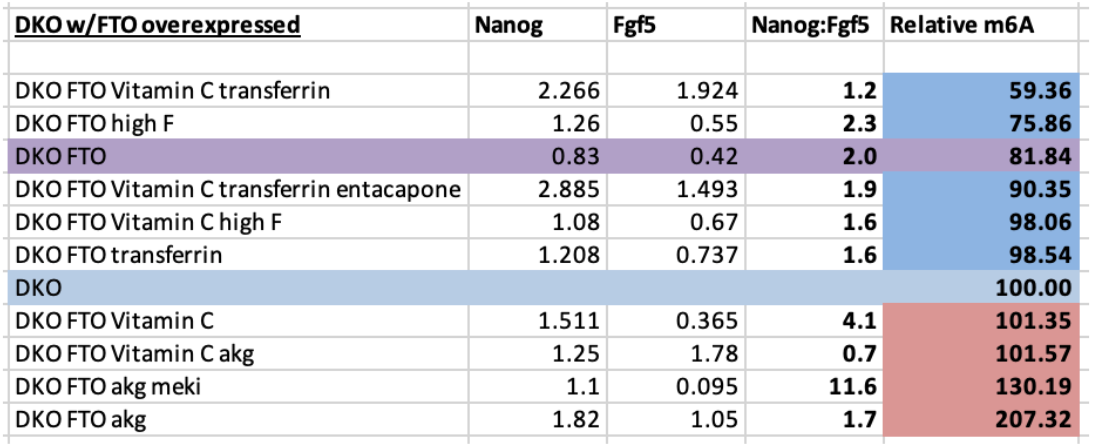

Since Vitamin C and transferrin could also be affecting the function of other demethylase enzymes, such as Jumonji-domain (histone demethylation) and Tet (DNA demethylation) proteins, we tested whether the FTO inhibitor entacapone [32] would mitigate the effects of Vitamin C and transferrin. Indeed, *Gsk-3* DKO ESCs treated with Vitamin C, transferrin and entacapone now showed a mere 9.6% reduction in m^6^A, essentially reversing the effects of Vitamin C and transferrin.

### Conclusions

Several of the small molecules that have been reported to promote retention of ESC pluripotency – alpha-ketoglutarate, Vitamin C, Gsk-3 inhibitors – also regulate the activity of the RNA demethylase, FTO. We sought to determine if there was a mechanistic connection between pluripotency and RNA methylation beyond what has already been described. More specifically, we wanted to learn if the changes in FTO activity were one of the main drivers of pluripotency and not merely a downstream consequence. While our data does not resolve this question, we did find an intriguing and potentially practical discovery – that the combination of Vitamin C and transferrin was a potent effector of FTO activity, driving m^6^A levels well below that seen with any other small molecule combination. FTO is a 2-oxoglutarate-dependent dioxygenase enzyme, and has been shown to require reduced iron (Fe^+2^) for its activity [34]. Iron is present in DMEM and likely in fetal bovine serum as well. Enhanced transport of iron into cells would facilitate its binding to FTO. Since transferrin naturally binds and transports iron into cells, the addition of transferrin to media should enhance iron transport. However, the iron needs to be reduced from Fe^+3^/Fe^+4^ to Fe^+2^. This is where Vitamin C is critical, acting as the reducing agent to produce the Fe^+2^ required for FTO function. This paradigm demonstrates why neither Vitamin C nor transferrin alone was sufficient to induce the potent reduction in m^6^A seen when the two molecules were combined. Practically speaking, we believe that the addition of Vitamin C and transferrin to ESC media would greatly benefit research labs who routinely culture mouse ESCs. It will be interesting to determine if the same effect on FTO activity and m^6^A levels is seen in human ESCs and iPSCs. One possible hint comes from a study showing that Vitamin C and Gsk-3 inhibitors potently promoted iPSC derivation when combined with Yamanaka factors as opposed to OSKM alone [7]. Since Gsk-3 inhibition increases FTO protein levels and Vitamin C reduces the iron required for its function, it seems plausible that the effects of Vitamin C and Gsk-3 inhibitors are to affect m^6^A levels. It would also be interesting to see if the addition of transferrin enhanced iPSC derivation even further.

## Materials and Methods

### Cell Culture and Transfection

Low passage, feeder-free wild-type (WT) mouse (E14K) and *Gsk-3* DKO ESCs, were grown on 0.1% gelatin-coated plates with standard DMEM containing 4.5 g/L glucose (Gibco) supplemented with 15% fetal bovine serum (HyClone), 1% non-essential amino acids (Gibco), 1% sodium pyruvate (Gibco), 1% L-glutamine (Gibco), 1% penicillin/streptomycin (Gibco) 55 μM 2-mercaptoethanol (Gibco), and 1000 units/mL recombinant leukemia inhibitory factor (LIF). Media was replenished every other day. The following small molecules were dissolved in sterile water and added to media at the following final concentrations: L-ascorbic acid 2-phosphate trisodium salt (Wako), 50 mg/mL; recombinant human transferrin (InVitria), 80 μg/mL; dimethyl 2-oxoglutarate (Sigma Aldrich), 3mM; NADH disodium salt (EMD Millipore), 300 μM; NADPH tetrasodium salt (EMD Millipore), 300 μM; PD0325901 (Selleckchem), 10 μM; SB415286 (Tocris), 30 μM; and entacapone (Selleckchem), 25 μM.

WT and *Gsk-3* DKO ESCs were transfected using PEI [33]. 1 x 10^6^ ESCs/well were resuspended in OptiMEM (Gibco) with 1800 ng of pCAGEN-FLAG-FTO, along with 200 ng of pMax-GFP. After incubation for 30 min at room temperature, cells were added to gelatin-coated 6-well plates containing complete media. Successful transfections were confirmed after 18 hours via fluorescence microscopy (Evos FL; ThermoFisher Scientific). When treating transfected cells with inhibitors, media was removed after 18 hours and replaced with fresh media containing small molecules. Cells were then harvested 24 hours later.

### RNA Isolation, cDNA Synthesis and Quantitative PCR

1 x 10^6^ ESCs/well were plated on gelatin-coated 6-well plate. RNA was isolated using TRIzol reagent (ThermoFisher Scientific) and extracted using Direct-zol RNA Miniprep columns (Zymo) following the manufacturer’s protocol. RNA was quantified using a Nanodrop One^C^ (ThermoFisher Scientific). 2 μg of total RNA was used to synthesize cDNA using the High Capacity Reverse Transcriptase kit (Applied Biosystems) according to the manufacturer’s protocol. The amount of input RNA used was kept constant for each RT reaction. Reactions were run on a StepOne Real-Time PCR System (Applied Biosystems) using PrimeTime gene expression master mix (IDT) and PrimeTime qPCR assays, *Nanog* (Mm.PT.58.23510265) and *Fgf5* (Mm.PT.58.17838080) (IDT). Three biological replicates and three technical replicates were used for WT and *Gsk-3* DKO ESCs. All threshold cycle (Ct) values were normalized to a mouse *Gapdh* endogenous control (Mm.PT.39a.1) (IDT), and relative quantification was calculated from the median Ct value.

### m^6^A Quantification

m^6^A levels were quantified using the EpiQuik m^6^A RNA Methylation Quantification Kit (Epigentek) following the manufacturer’s instructions. Briefly, 200 ng of total RNA from ESCs was bound to wells in a clear 96-well plate via RNA high binding solution, then incubated at 37°C for 90 minutes. Capture antibody was incubated at RT for 60 minutes, washed 3x, incubated with detection antibody for 30 minutes at RT, washed 3x, and finally incubated with enhancer solution for 30 minutes at RT. After washing 5x, colorimetric detection of the signal was obtained by incubating with a developer solution and then a stop solution, followed by detection at OD 450 nm using an EnSpire plate reader (PerkinElmer). A standard curve of m^6^A RNA (provided with kit) was included on each plate, in duplicate, ranging from 0.02 ng to 1 ng. The OD readings were then used to calculate the relative quantification using the following formula:

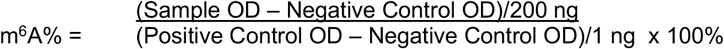

## Abbreviations

m^6^A: methyladenosine
ESCs: embryonic stem cells
Gsk-3: glycogen synthase kinase-3
DKO: double knockout
LIF: leukemia inhibitory factor
Fgf5: fibroblast growth factor 5

## Acknowledgements

This research is supported by NIH R15GM119103 (C.J.P.)

